# Mitochondria transported by Kinesin 3 prevent localized calcium spiking to inhibit caspase-dependent specialized cell death

**DOI:** 10.1101/2025.01.08.631976

**Authors:** Rashna Sharmin, Aladin Elkhalil, Sara Pena, Pranya Gaddipati, Ginger Clark, Pavak K. Shah, Mark W. Pellegrino, Shai Shaham, Piya Ghose

## Abstract

Polarized cells (such as neurons), defined by distinct compartments are, like many cell types, subject to developmental elimination, as in neurite pruning. The molecular mechanism behind specialized elimination remains a largely open question. We previously introduced the “tri-partite” embryonic cell death program Compartmentalized Cell Elimination (CCE) in the scaffolding tail-spike epithelial cell and sex-specific CEM neurons of *C. elegans*. CCE is stereotyped and ordered, with three distinct programs eliminating three cell compartments, and bearing morphological hallmarks reminiscent of neurite pruning. Here we report first, that, prior to CCE onset, mitochondria undergo UNC-116/Kinesin 1-dependent irreversible retrograde transport; and second, that the caspase protease CED-3 promotes the completion of the proximal nicking event of CCE by helping inhibit the kinesin 3 homolog UNC-104. While known canonically to carry only synaptic vesicles, UNC-104/Kinesin 3, in the CCE context, and in the absence of CED-3/Caspase, can transport mitochondria anterogradely to the severing site. We observe both caspase activity and Ca^2+^ spiking at this site prior to nicking. Mitochondria appear to protect against nicking in the absence of CED-3 via their MCU-1 uniporter-dependent Ca^2+^ uptake capacity. Our study sheds light on the molecular machinery of specialized cell elimination and pruning, highlighting involvement of region-specific Ca^2+^ signaling, an anti-death function of mitochondria via local Ca^2+^ uptake, mitochondrial transport as a regulatory strategy, with UNC-116/kinesin 1-mediated retrograde transport important for cell elimination priming and UNC-104/kinesin 3 in a previously undescribed role as a non-canonical anterograde mitochondrial motor.

## Introduction

Programmed cell death is a vital feature of normal development and homeostasis(1), with the programmed elimination of neurons a critical facet of neurodevelopment(2). A large proportion of neuronal connections in the early brain are supernumerary and are developmentally eliminated(3) through the removal of entire cells or region-specifically(4). Region-specific neuronal elimination may be in the form of both neurite and synaptic pruning(3, 5). Failure in programmed neuronal regression can result in numerous neurodevelopmental and psychiatric disease conditions(6, 7). Programmed neuronal death and localized neuronal elimination show morphological overlaps with neurodegeneration under pathological conditions or following injury(8). As such, a fundamental understanding of developmental pruning and death of neurons is of therapeutic significance. The molecular mechanisms of compartment-specific regressive events broadly are still being elucidated.

The morphological complexity of neurons makes them an especially intriguing setting to study cell elimination. Intricate and highly polarized, neurons are the most elaborate cells of the body, with structurally and functionally distinct domains, which is a major consideration when it comes to eliminating these kinds of cells. Signaling in such morphologically complex cells at large is inherently compartmentalized, requiring the integration of information between and across compartments. Such compartments can be very distinct both structurally, molecularly and functionally each with different contributions to cell function and fate. For example, axons and dendrites have vastly different roles in execution of neuronal communication. Also, the axon initial segment (AIS), a specialized subcellular domain that precisely demarcates neuronal compartments at (9, 10) the proximal axon, is important for neuronal excitability, maintaining neuronal polarity(11, 12) and for the protein trafficking and distribution across the somatodendritic and axonal compartments (13).

Region specificity of functions can be accomplished via differential signaling events. For example, local Ca^2+^ signaling occurs in compartments such as such as presynaptic active zones and postsynaptic densities(13), or local energy production by mitochondria directed to energy-demanding regions, such as synapse or injury sites(14, 15). The compartments of neurons and other polarized cells also communicate with each other(16). One important way to achieve inter-compartmental communication is through intracellular transport via motors that carry various cargo along the cytoskeleton, as in microtubule motors such as kinesins and dyneins(17, 18). Cargo can include synaptic vesicles for neurons, and organelles(19), such as lysosomes and mitochondria, expanding to other cell types as well. Whether organellar transport can promote local cellular regression is an open question. Additionally, different motors have different characteristics and are equipped to carry specific cargos and can be regulated by other adaptor molecules(20), post-translational modifications(21), and can be directed in response to local cellular cues, such as neuronal activity(22). How cargo specificity and directionally of motors is regulated is another long-interesting question.

The compartmentalized nature of specialized cells, thus, is a major consideration in understanding how such cells are eliminated. This is not limited to neurons but is also true for non-neuronal cells. Epithelial cells are also polarized into distinct structural and functional domains and epithelial cell death is commonplace during development and tissue homeostasis(23). For example, cell death is a feature of the gut epithelium during inflammation. How inter-compartmental communication, intracellular organellar transport, and local signaling events may cooperate to execute a specialized cell elimination program is a fascinating area to explore.

Of known cell elimination programs, apoptosis is the best characterized, with defined morphological hallmarks and genetics, with the core components known to be conserved across species(1) and relevant for disease such as in cancer therapeutics(24). The genetic regulation of apoptosis was first delineated in the nematode *Caenorhabditis elegans*(25, 26). A cardinal molecular feature of apoptotic cell death is its regulation by cysteine proteases, caspases(27, 28). Caspases have both apoptotic and non-apoptotic functions(27, 29, 30). Many caspase substrates have been implicated by *in vitro* through proteomic studies(31), though few caspase-regulated factors have been validated *in vivo*(32, 33). Several forms of non-apoptotic, caspase independent cell death modalities have also been described(34) and caspases have also been shown to play a localized role in developmental pruning(3).

We sought to explore how the fundamental feature of polarized cells, compartmentalization, affects the elimination of this class of cells. We adopted an unbiased forward genetic approach in the nematode *C. elegans* to explore these fundamental principles of complex cell elimination and to identify novel regulators. We have previously introduced the embryonic cell death phenomenon, Compartmentalized Cell Elimination (CCE)(35) taking place in both the embryonic tail-spike epithelial cell (TSC) (**FIG 1A**) and sex-specific CEM sensory neurons of *C. elegans*. Briefly, CCE is a tripartite developmental cell death program with hallmarks of developmental neurite pruning. Using TSC-specific membrane markers, TSCp::myrGFP(35) and TSCp::myrmKate2 (**FIG 1A**) we found that the first visible hallmark of CCE is the start of a severing or nicking event at the proximal segment of the TSC process near the cell body (**FIG 1B**). Following this, the cell soma rounds, and the proximal segment of the process undergoes beading followed by fragmentation (**FIG 1C**), and the distal segment retracts into a distal node(35)(**FIG 1D**). The proximal process segment is removed fully first followed by the stochastic phagocytosis of the soma and distal remnants by different phagocytes. CCE, while dependent on the main *C. elegans* caspase CED-3, is genetically distinct from apoptosis(36) and has led to the identification of novel regulators of programmed cell death(35–38).

**Figure 1.**
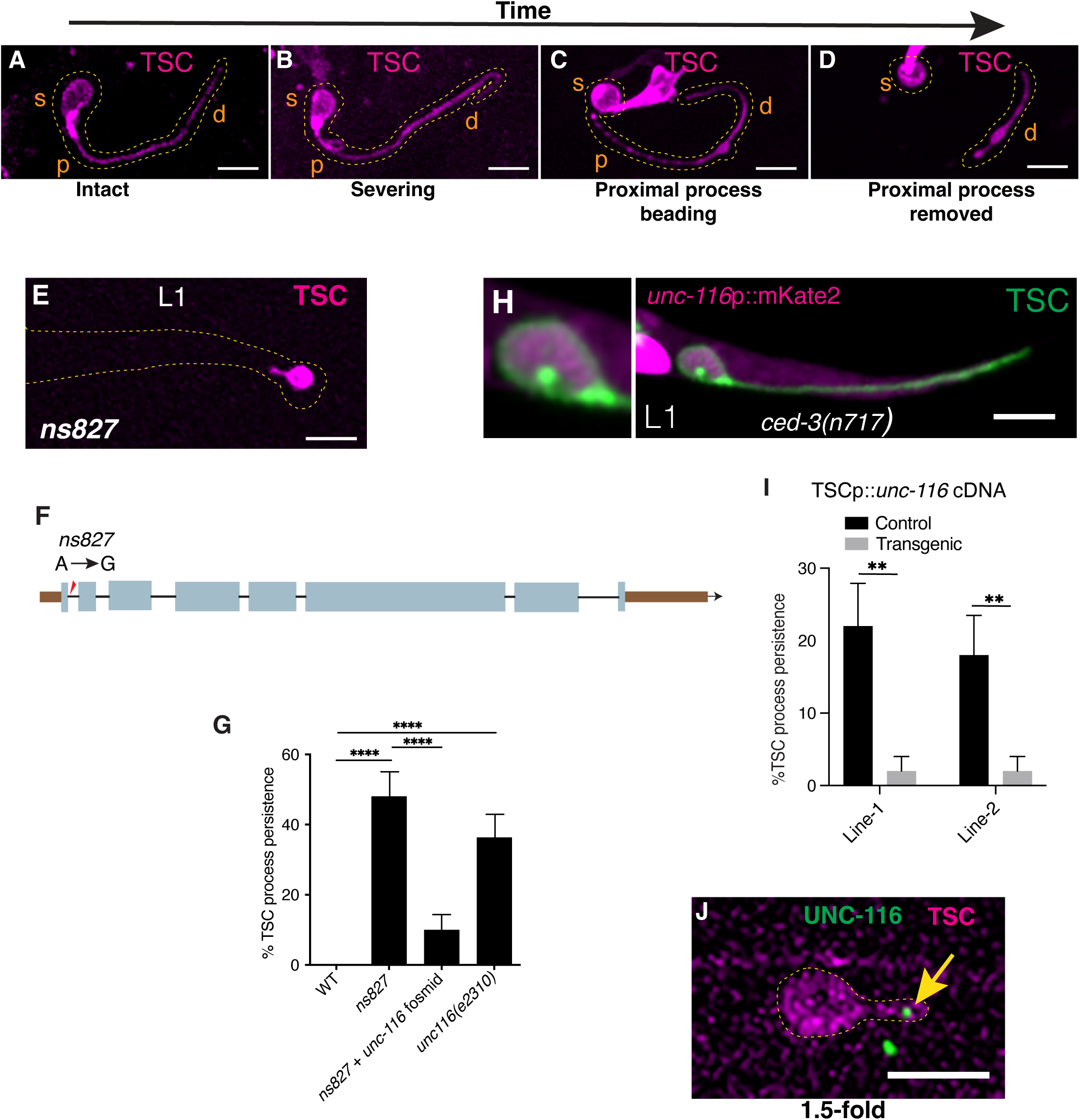
UNC-116/Kinesin 1 promotes CCE. (A-D) Steps of CCE. s, soma; p proximal process; d, distal process. **(E)** L1 *unc-116 (ns827))* mutant. n=50 biologically independent animals with similar results. **(F)** *unc-116* gene structure**. (G)** Process elimination defects in indicated genotypes. n=sample sizes for statistics for each bar=50, with n referring to number of biologically independent animals. **(H)** Expression of *unc-116* as seen using a transcriptional reporter for *unc-116* in TSC. n=20 biologically independent animals with similar results. **(I)** *unc-116 (ns827)* mutant TSC-specific rescue. n=sample sizes for statistics=50 with n referring to number of biologically independent animals. **(J)** Endogenously tagged (GFP, green) UNC-116/Kinesin 1 showing localization in TSC distal process (magenta) at the 1.5-fold embryo stage. n=20 biologically independent animals with similar results. Scale bar, 5μm.

Here we identify two opposing microtubule motors, UNC-116/Kinesin 1 and UNC-104/Kinesin 3 as regulators of CCE. We report that irreversible retrograde mitochondrial transport via UNC-116/Kinesin 1 from the cell process/dendrite is important for CCE priming, prior to visible cell death onset. We also report mitochondria as a non-canonical cargo for UNC-104/Kinesin 3, which is conventionally associated with synaptic vesicle transport(13), and that short range anterograde transport of mitochondria can protect against CCE. We report mitochondria as cytoprotective through their Ca^2+^ uptake ability via the Ca^2+^ uniporter MCU-1. We implicate CED-3/Caspase in the negative regulation of UNC-104/Kinesin 3 and region-specific caspase activity, and local Ca^2+^ spiking as essential for CCE to fully progress. Collectively, our findings implicate mitochondria and new contexts for their transport as regulators of neurite pruning and possibly, by extension, neurodegeneration in disease or injury contexts, and shed new light on the genetic regulation of complex cell elimination.

## Results

### UNC-116/Kinesin 1 is required for proper CCE

We previously performed a forward genetic screen seeking to uncover factors regulating CCE, employing a membrane marker labelling the TSC (TSC-myrGFP)(35). From this screen, we recovered a mutant, *ns827,* in which the distal segment of the TSC process persists in first larval stage (L1) animals, when the TSC would normally be absent following its embryonic elimination (**FIG 1E**). Following Whole Genome Sequencing, Single Nucleotide Polymorphism (SNP) mapping and genomic DNA rescue, we identified the causative lesion to be an A to G splice donor after exon 1 of the gene *unc-116* (**FIG 1F, G**). The gene *unc-116* encodes the nematode homolog of the mammalian microtubule plus-end directed motor KIF5/Kinesin 1(39). We found *unc-116* to be expressed in the TSC by examining a transcriptional reporter for *unc-116* (*unc-116*pro::mKate2) (**FIG 1H**). We next performed cell-specific rescue experiments and found that UNC-116/Kinesin 1 functions in the TSC cell-autonomously (**FIG 1I**). We also looked at the localization of UNC-116/Kinesin 1 endogenously tagged with GFP and found UNC-116/Kinesin 1 to be present in the tail-spike cell and localized at the distal tip of the process at the 1.5-fold embryo stage (**FIG 1J**).

### UNC-116/Kinesin 1 dependent irreversible retrograde mitochondrial transport primes CCE

We proceeded to inquire about the identity of UNC-116/Kinesin 1’s cargo and function in the TSC. Based on the mutant phenotype, we posited that in the absence of this motor, cargo is restrained in the distal process, preventing this cell region from elimination and that UNC-116/Kinesin 1 functions in retrograde transport of this cargo. We also speculated that the cargo has region-specific cytoprotective capability. Kinesin 1 is a conserved motor well-known to carry mitochondria. Prior studies in *C. elegans* show UNC-116/Kinesin 1-dependent mitochondria to be important for neurite preservation following injury(40). Mitochondria have long been associated with both cell survival and destruction(41). We predicted that mitochondria may be the relevant UNC-116/Kinesin 1 retrograde cargo for CCE with a compartment-specific pro-survival capability. To test this, we generated a reporter for mitochondria (TSC-specific Mito-Matrix-GFP) against a red membrane marker to visualize mitochondria in the TSC in wild type animals. We first used this reporter to test if mitochondria are present in the remaining process of the *unc-116* mutants. We indeed found mitochondria present in the persisting TSC process remnant consistently (30/30 animals) (**FIG 2A-C**).

**Figure 2.**
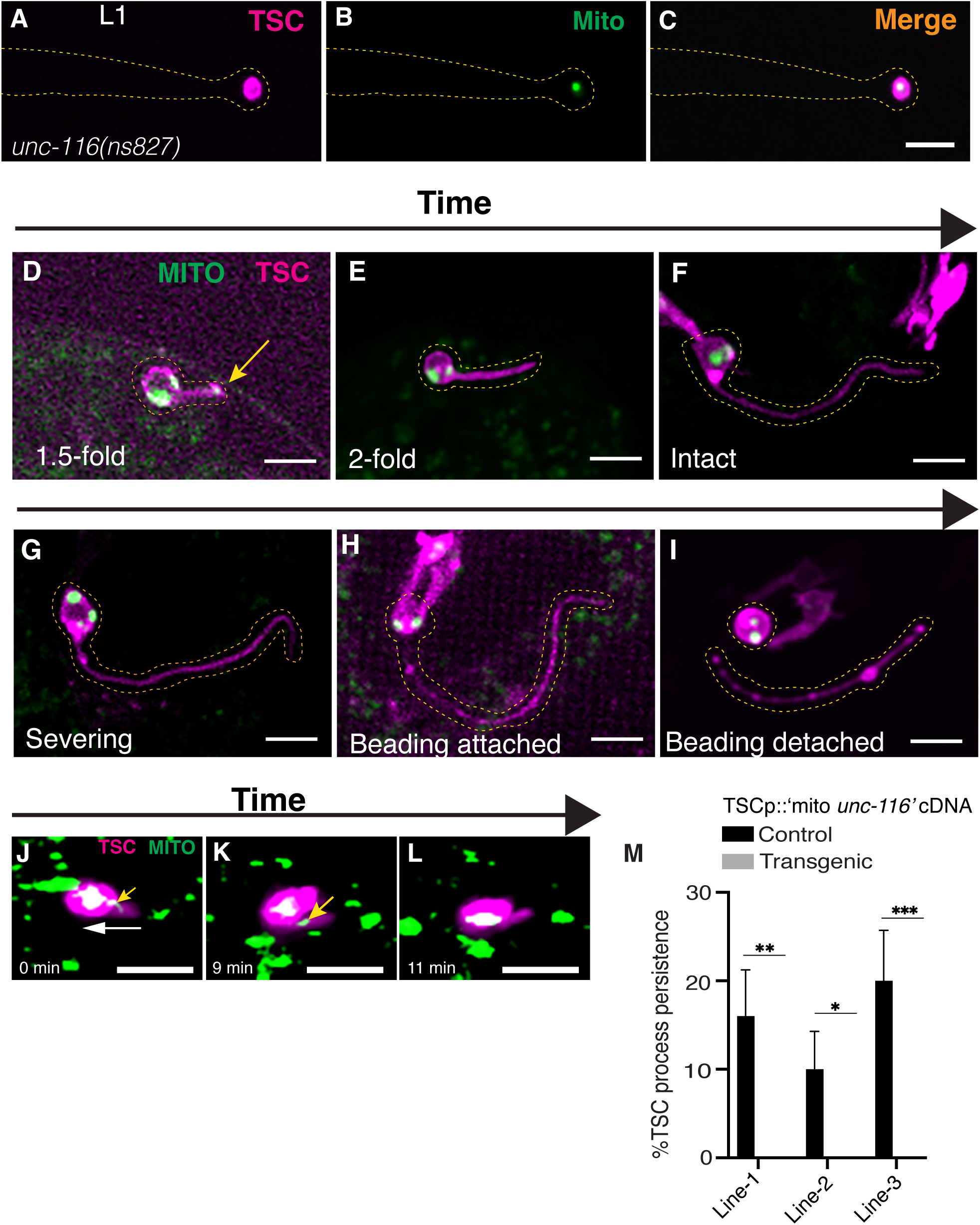
Irreversible retrograde transport of mitochondria via UNC-116/Kinesin 1 precedes CCE. (A-C) Image showing persisting mitochondria in L1 *unc-116 (ns827)*. n=20 biologically independent animals with similar results. **(D-I)** Mitochondria position across CCE stages in TSC in still images. n=10 biologically independent animals with similar results. **(J-L)** Frames from time-lapse movie **(Supplemental Movie 1)** of mitochondrial transport in TSC. n=2 biologically independent animals with similar results. **(M)** Mitochondria-specific rescue of *unc-116* mutant CCE defect. n=sample sizes for statistics for each bar=50 with n referring to number of biologically independent animals. Scale bar, 5μm.

We next examined TSC-Mito::GFP in wild-type embryos first as still images at specific timepoints and found mitochondria were only seen in the process distally at or before the 1.5-fold embryo stage (**FIG 2D-I**), noting UNC-116/Kinesin 1 is also clearly localized distally at this early stage (**FIG1 J**). To test the idea that mitochondria are transported into the soma, we performed time lapse imaging using a light-sheet microscope (**FIG 2J-L, Supplemental Movie 1**) and found mitochondria to move retrogradely, entering the soma.

We next performed a rescue experiment using “TSCp::Mito-UNC-116” in which UNC-116/Kinesin 1 is linked to the outer membrane protein Tom 7 such that this motor can only transport mitochondria as cargo, based on a previously used design(40) (**FIG 2M**). We saw strong rescue with “TSCp::Mito-UNC-116” suggesting that transport of mitochondria by UNC-116/Kinesin 1 can specifically allow for proper CCE.

### In absence of localized caspase activity mitochondria appear at the severing site and CCE reverses

We found from time-lapse imaging of wild-type animals harboring TSC-Mito::GFP (**FIG 3A-D’, Supplemental Movie 2**) that mitochondria do not appear at the soma-process junction at the start of CCE, where the proximal severing/nicking event of CCE occurs (**FIG 3B, B’**). We asked about a possible correlation between this severing event and mitochondrial localization and turned to the genetics of CCE to assess the same. The main *C. elegans* caspase CED-3 is essential for CCE, though it is regulated in a non-canonical manner, independent of EGL-1/BH3-only of the canonical apoptotic pathway(36). In the absence of CED-3, TSC persists intact in larva(35, 36). Unexpectedly, in *ced-3* mutants, most CCE hallmarks do appear, including soma rounding, proximal process beading and distal node/varicosity formation (**FIG 3E-H’, Supplemental Movie 3**). However, the severing event seen in wild-type embryos remains incomplete (**FIG 3F, F’**), and even more surprising, the cell appears to revert to its original morphology (**FIG 3H, H’**). This suggests the severing event is required for CCE to proceed properly to completion and that CED-3/Caspase is important for this step. To examine region-specificity, we tested a GFP::C3Ai caspase activity reporter(42) we expressed in the TSC in wild-type (**FIG 3I-L**) and *ced-3(n717)* mutants (**FIG 3M-P**). We observed a sharp and specific signal from this reporter at the soma-process junction in wild-type (**FIG 3I, J**), but not in *ced-3(n717),* suggesting that CED-3 caspase activity is at play at the severing site as CCE sets in.

**Figure 3.**
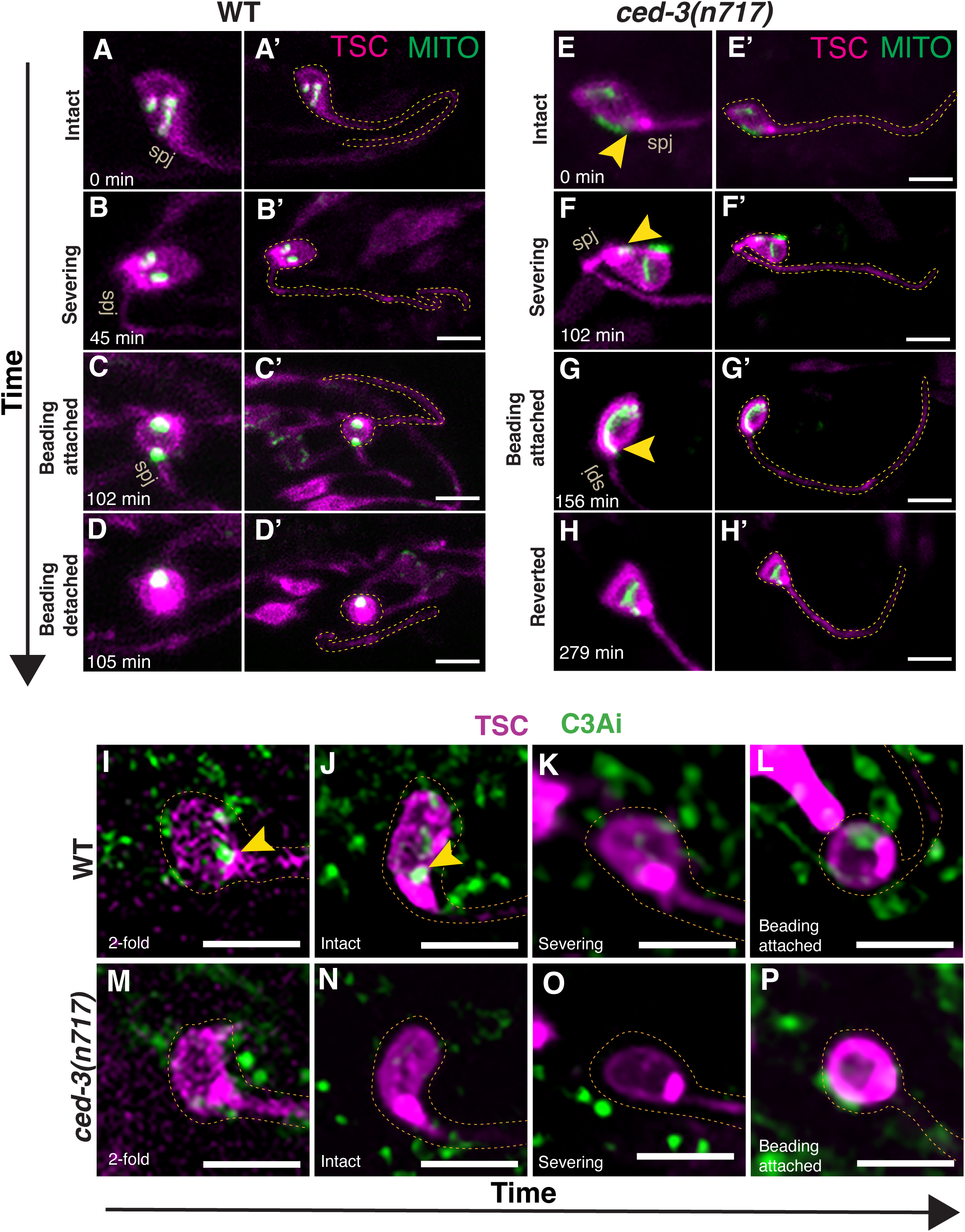
In absence of localized caspase activity, mitochondria appear at the severing site and CCE reverses. Stills from movie showing mitochondrial movement during CCE in wild-type **(A-D’) (Supplemental Movie 2)**. and *ced-3(n717)* (E-H’) (Supplemental Movie 3) embryos, showing mitochondria entering soma process junction anterogradely in *ced-3(n717)* mutants (**E-G’**), n=2 biologically independent animals with similar results. **(I-P)** Caspase activity reporter GFP::C3Ai dynamics across CCE stages in wild-type **(I-L)** and *ced-3(n717)* (M-P). n=20 biologically independent animals with similar results. Scale bar, 5μm.

Of note, unlike wild-type, in *ced-3* mutants, mitochondria are seen to travel to this region anterogradely from the soma in the embryo (**FIG 3G, H**). This led us to hypothesize that, the absence of CED-3, mitochondria are carried by an anterograde motor and can somehow functionally prevent severing and proceeded to explore this idea.

### UNC-104/Kinesin 3 acts as a mitochondrial anterograde motor negatively regulated by CED-3/caspase

As we have previously reported, the *ced-3(n2427)* hypomorph has a 40% CCE defect of inappropriate TSC persistence(35). We reasoned that if a double mutant in this background with any gene suppresses this defect, the protein encoded by that gene may be negatively regulated by CED-3/Caspase. We tested multiple genes in this genetic background. Curiously, we did not see changes when testing genes encoding known mitochondrial motors, *trak-1* and *dhc-1/dynein*, or *unc-116* (**FIG 4A**). However, unexpectedly, a double mutant for *ced-3(n2427); unc-104(e1265)* showed a significantly reduced CCE defect (**FIG 4B**) compared to the single mutant *ced-3(n2427)*, though the *unc-104* single mutant did not show a noticeable CCE defect alone. We reintroduced *unc-104* into the TSC specifically in *ced-3(n2427); unc-104(e1265)* and observed significant rescue (return to the *ced-3* hypomorph defect) (**FIG 4B**), suggesting autonomous function in the TSC. UNC-104 is the nematode homolog of mammalian Kinesin 3, another plus-end directed microtubule motor protein known for its role in synaptic vesicle transport.

We visualized UNC-104 endogenously in wild-type versus *ced-3(n717)* mutants by generating a CRISPR/Cas9 endogenously tagged (GFP at C terminus) *unc-104*. We did not observe UNC-104::GFP puncta in the TSC in in wild-type after the 1.5-fold embryo stage (**FIG 4C-F**), but could see signal in *ced-3(n717)* mutants at the soma-process junction (**FIG 4G-J**).This supports a model in which UNC-104/Kinesin 3 is stable in the absence of CED-3/Caspase, and for a role in the regulation of severing.

**Figure 4.**
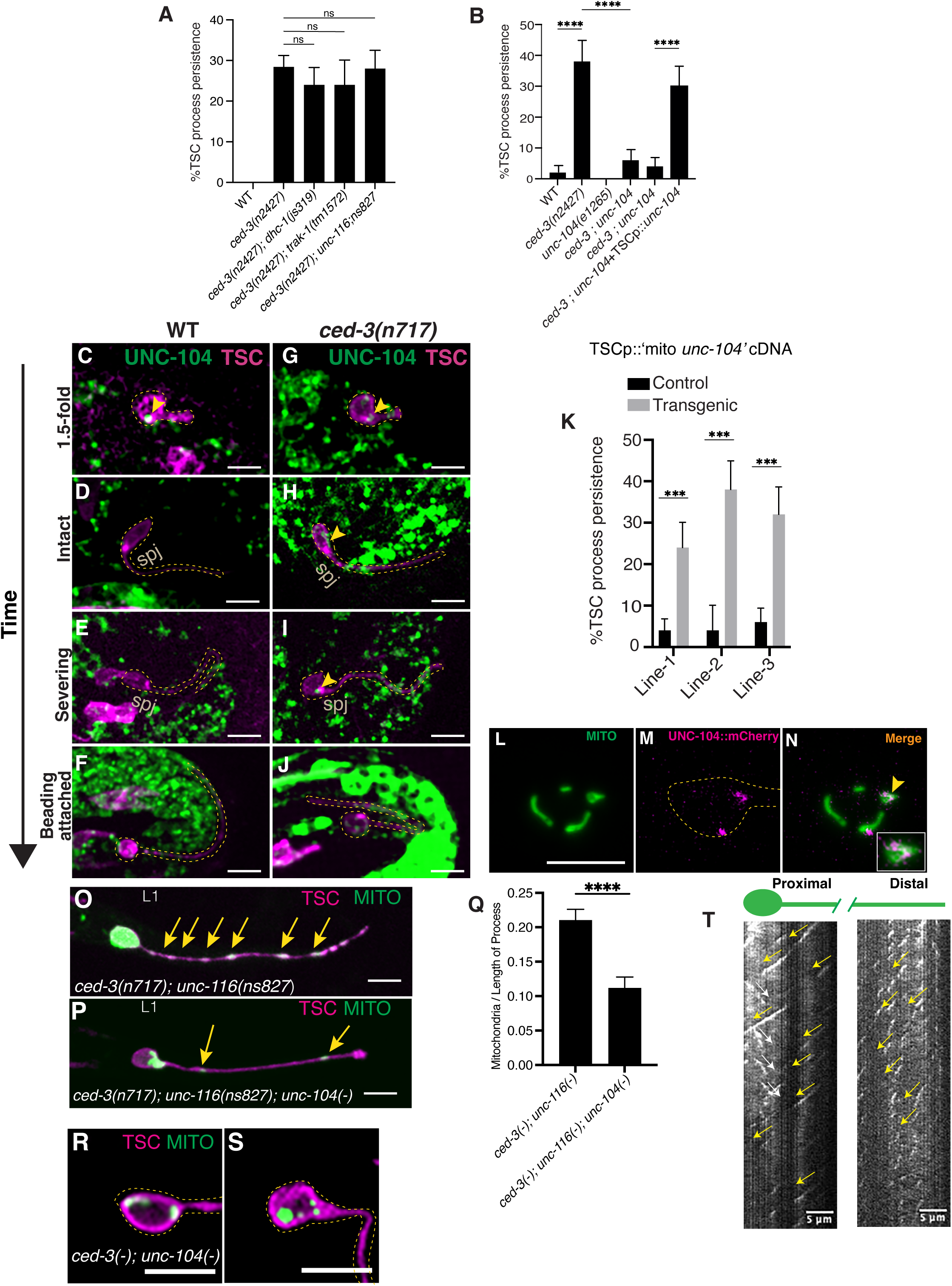
UNC-104/Kinesin 3 acts as a short range mitochondrial anterograde motor negative regulated by CED-3/Caspase. (A, **B)** Graphs for anterograde mitochondrial motor candidate gene screen. n=sample sizes for statistics for each bar=50 with n referring to number of biologically independent animals. **(C-F)** Lack of UNC-104 (endogenously tagged with GFP) signal in wild-type after 1.5-fold embryo stage. **(G-J)** Localization of UNC-104 endogenously tagged with GFP at different time points in *ced-3(n717)* embryos showing presence in TSC (magenta) after the 1.5-fold embryo stage. n=10 biologically independent animals with similar results. **(K)** TSC mitochondria-specific rescue of *unc-104* mutant CCE defect. n=sample sizes for statistics for each bar=50 with n referring to number of biologically independent animals. **(L-N)** Co-localization of UNC-104 and mitochondria in L1 larva of *ced-3(n717)*. n=10 biologically independent animals with similar results. **(O)** Enrichment of mitochondria in TSC process of *ced-3(n717)*; *unc-116* double mutants. **(P)** Loss of this enrichment of mitochondria in TSC process of *ced-3(n717)*; *unc-116; unc-104* triple mutants. **(Q)** Graph for (P) and (Q), n=sample sizes for statistics for each bar=50 with n referring to number of biologically independent animals. **(R, S)** Mitochondrial localization in *unc-104(e1265); ced-3(n717)* mutants during CCE showing mitochondria absent at the soma-process junction. R, TSC still intact; S, TSC reverted from CCE after severing attempt. **(T)** Kymograph of TSC-EBP-2::GFP of *ced-4(n1162)* mutant L1 larvae **(Supplemental Movie 4 and 5).** n=10 biologically independent animals with similar results. Scale bar, 5μm.

We introduced UNC-104/Kinesin 3 linked to mitochondria TSCp::Mito-UNC-104 into *ced-3(n2427); unc-104(e1265)* with a red TSC membrane marker and we found significant rescue of the double mutant phenotype (**FIG 4K**), supporting the unexpected role of UNC-104/Kinesin 3 in mitochondrial transport that may directly or indirectly be inhibited by in a CED-3/Caspase. Of note, mitochondria are not known to be a classical cargo of UNC-104, which, rather, is associated with vesicle transport(13). We then tested for colocalization between a UNC-104/Kinesin 3, tagged with mCherry, and MitoGFP. We examined for colocalization (**FIG 4L-N**) in *ced-3* mutant L1 larval animals, in which we propose UNC-104/Kinesin 3 can remain stable, and found co-localization specifically at the soma-process junction.

Next, to test whether UNC-104/Kinesin 3 carries mitochondria anterogradely, we looked at an *unc-116(ns827)* (in which retrograde transport fails) as a double with *ced-3(n717)* (in which UNC-104/Kinesin 3 is presumed to be stable) (**FIG 4O**). Mitochondria were heavily enriched in the process in this background. We reasoned that this is due to the combination of a failure of retrograde mitochondria transport and excess anterograde transport. We next compared this to a triple mutant for *ced-3(n717)*; *unc-104(e1265); unc-116(ns827*) (**FIG 4P**) and found mitochondria to be in significantly lower number in the process of the intact cell in the absence of UNC-104/Kinesin 3 (**FIG 4Q**), suggesting UNC-104/Kinesin 3 is necessary for the enhanced mitochondrial presence in the double mutant, implicating it as an anterograde motor for mitochondria in the TSC that functions in absence of CED-3/Caspase. We also examined *ced-3(n717)*; *unc-104(e1265)* double mutant embryos during CCE and observed that, unlike *ced-3(n717)* single mutants, in which mitochondria appear at the soma-process junction (**FIG 3E-F**), mitochondria are absent at this site (**FIG 4R, S**), appearing more as wild-type (**FIG 3A-D**’). This supports the idea that UNC-104/Kinesin 3 can transport mitochondria to the soma-process junction in the absence of CED-3/Caspase.

### UNC-116 and UNC-104 play opposing roles in mitochondrial transport to regulate CCE

Our data suggest that UNC-116/Kinesin 1 and UNC-104/Kinesin 3 regulate CCE and mitochondrial transport in opposite ways, with UNC-116/Kinesin 1 promoting CCE in a wild-type context and UNC-104/Kinesin 3 preventing it in the absence of CED-3/Caspase. UNC-116/Kinesin 1 removes mitochondria from the process through irreversible retrograde transport that promotes CCE. Conversely, UNC-104/Kinesin 3 can only function when CED-3/Caspase is absent and transports mitochondria anterogradely into the severing site to prevent CCE. Curiously, both UNC-116/Kinesin 1 and UNC-104/Kinesin 3 are plus-end directed motors yet carry mitochondria in specific and opposite directions.

For this model to hold, microtubule orientation in the TSC process must be of mixed polarity, both plus-end-in and plus-end-out. To test this, we generated a TSC-specific reporter for the microtubule plus end binding protein EBP-2(43). We observed TSC process EBP-2 comets directed both to and away from the TSC soma (**FIG 4T, Supplemental Movie 4 and 5**), supporting the idea that microtubules of the TSC process are of mixed polarity and reconciling how two plus-end directed proteins may regulate mitochondrial transport and death of the TSC. Interestingly, mixed microtubule polarity is a characteristic to dendrite architecture, and the CCE steps of the TSC process are seen in the single dendrite of the hermaphrodite CEM sensory neurons(35). This suggests the TSC process may serve as a viable model for dendrite elimination.

We then asked: if UNC-104/Kinesin 3 and UNC-116/Kinesin 1 are both plus-end directed motors and the TSC process is of mixed polarity, why would these motors be committed to transport mitochondria in a specific direction, retrogradely for UNC-116 and anterogradely for UNC-104? We reasoned that this may be due to where these proteins are originally localized as the TSC first becomes polarized. As mentioned, endogenously tagged versions of these motors against a TSC membrane mCherry marker revealed that UNC-116/Kinesin 1 is initially localized distally (**FIG 1J**) and UNC-104 more proximally, close to the soma (**FIG 4G-J**). This suggests the bidirectional mitochondrial movement may be due to the combination of opposite pole origins of the two plus-end directed motors and mixed microtubule polarity of the TSC process. We did note also that the mixed microtubule polarity is seen more proximally and the plus-end-in orientation distally (**FIG 4T**). The observation that the plus-end-out EBP-2 movement is restricted proximally may be related to the short-range anterograde nature of UNC-104.

### Mitochondria are protective via their Ca^2+^ uptake potential

Our data implicate mitochondria as protective against CCE and that their transport by UNC-116/Kinesin 1 and UNC-104/Kinesin 3 is important to regulate cell elimination. To examine the nature of this protective role, we first tested if mitochondrial energy production gives mitochondria the ability to preserve the cell. We tested *clk-1*, which encodes co-enzyme Q, a component of the oxidative phosphorylation pathway(44), against the *ced-3(n2427)* hypomorphic background (**FIG 5A**). Surprisingly, the double mutant had the same proportion of CCE-defective animals as the *ced-3* hypomorph, suggesting mitochondrial energy production may not explain why mitochondria need to be removed from the process for its elimination. We next tested an alternative hypothesis that the ability of mitochondria to act as Ca^2+^ sinks may confer upon them protective capability, promoting cell survival. We therefore tested a mutant for the gene encoding the mitochondrial Ca^2+^ uniporter (MCU), MCU-1, a Ca^2+^-selective mitochondrial ion channel (45) in the *ced-3* hypomorphic background. Interestingly, in this instance, we did see suppression of the *ced-3* hypomorphic CCE defect (**FIG 5B**), suggesting the ability of mitochondria to import cytosolic Ca^2+^ of the TSC may give it a protective role. Supporting this, when we re-introduced *mcu-1* in the TSC specifically, we found the phenotype to be rescued (**FIG 5C**).

Given that (i) based on genetics, mitochondrial Ca^2+^ uptake appears to promote cell survival and prevent CCE, and that (ii) in the absence of CED-3/Caspase, UNC-104/Kinesin 3 and mitochondria localize to the soma-process junction prior to cell severing, we tested the hypothesis that region-targeted transport of mitochondria that take up Ca^2+^ protects the cell from elimination. This would suggest that a local increase in cytosolic Ca^2+^ is important for CCE. To test this, we examined a TSC-specific cytosolic GCaMP5a(46) reporter to visualize Ca^2+^ spikes in the cell. Interestingly, in wild-type embryos we detected cytosolic Ca^2+^spikes just before the soma-process junction severing stage (**FIG 5D-F**). However, when we then looked at *ced-3* mutants (**FIG 5G-I**) this severing point signal was absent. Given our observation that mitochondria and UNC-104/Kinesin 3 are localized to this region prior to severing, and that mitochondria appear to prevent CCE owing to their uptake of cytosolic Ca^2+^, we looked at our GCaMP5a reporter in both the *ced-3*(*-*)*; unc-104*(*-*) double mutants (**FIG 5J-L**) and *ced-3*(*-*)*; mcu-1*(*-*) double mutants (**FIG 5M-O**). As mentioned above (**FIG 4R, S**), mitochondria are not present at the severing point in *ced-3(n717); unc-104(e1265)* double mutants. As such, cytosolic Ca^2+^ would not be taken up and a GCaMP signal would be seen. In the latter case, mitochondria would be transported and present at the severing point, but mitochondrial Ca^2+^ uptake would not occur due to lack of MCU-1. In both cases, we would predict return of the GCaMP signal, even in the absence of CED-3/Caspase. Indeed, we found that the severing point GCaMP signal returned in both double mutants this solidifying an involvement of MCU-1 and UNC-104 in Ca^2+^ regulation and severing. We also noted that, despite the lack of Ca^2+^ uptake at the severing point, the TSC still persists in *ced-3*(*-*)*; unc-104*(*-*) double mutants and *ced-3*(*-*)*; mcu-1*(*-*) double mutants, though in *ced-3*(*-*)*; mcu-1*(*-*) double mutants, the persisting cell after hatching appears in a stage of degeneration (data not shown). Loss of MCU-1 and loss of UNC-104/Kinesin 3 would conceivably have different effects on the cell, with one impacting mitochondrial Ca^2+^ uptake broadly and another only affecting their short-range transport to a particular cell region. We do note that CED-3/Caspase likely has many downstream targets and related proteins, such that severing, while important for CCE, may be one of several steps needed for a complex process such as CCE. Indeed, caspases can have multiple substrates relevant to in cell death (31), with even non-killing roles(30). CED-3/Caspase could directly be a regulator of UNC-104/Kinesin 3.

**Figure 5.**
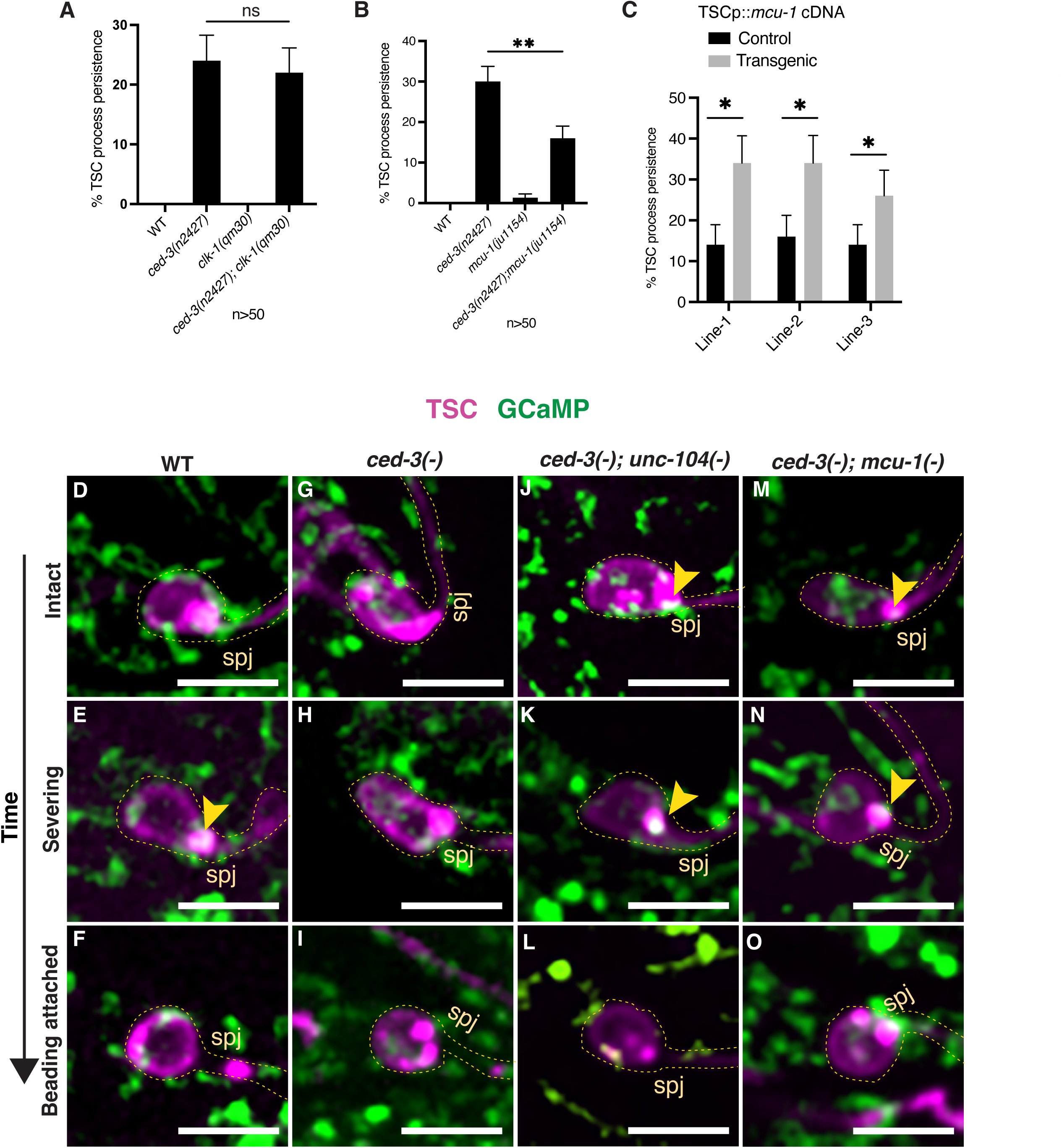
Mitochondria are protective via their Ca^2+^ uptake potential. **(A)** Graph for showing CLK-1 does not regulate CCE. n=sample sizes for statistics for each bar=50 with n referring to number of biologically independent animals. **(B)** Graph showing MCU-1/MCU prevents CCE. n=sample sizes for statistics for each bar=50 with n referring to number of biologically independent animals. **(C)**Graph showing MCU-1 functions in the TSC cell-autonomously to repress CCE. n=sample sizes for statistics=50 with n referring to number of biologically independent animals. Localization of GCaMP5 Ca^2+^ sensor in TSC across CCE stages in wild-type **(D-F)**, *ced-3(n717)* **(G-I)**, *ced-3 (n717); unc-104 (e1265)* double mutant **(J-L)** and *ced-3(n717); mcu-1(ju1154)* double mutant **(M-O).** n=10 biologically independent animals with similar results. Scale bar, 5μm.

In sum, here we implicate mitochondria as an anti-death organelle by virtue of its Ca^2+^ uptake capability, and cytosolic Ca^2+^ as a pro-death signaling factor the specific role of which in promoting CCE remains to be determined. We propose that CCE priming is ensured distally through irreversible retrograde transport of mitochondria into the soma by UNC-116/Kinesin 1. Afterwards, enhanced local cytosolic Ca^2+^ promotes the process-severing event of cell dismantling. Mitochondria must be kept remote from the severing point via CED-3/Caspase’s negative regulation of UNC-104/Kinesin 3. In the absence of CED-3/Caspase, mitochondria can be transported short-range via stable UNC-104/Kinesin 3 and protect the cell from severing and death, and this protective capacity of mitochondria depends on MCU-1-dependent Ca^2+^ uptake ability (**FIG 6**).

**Figure 6.**
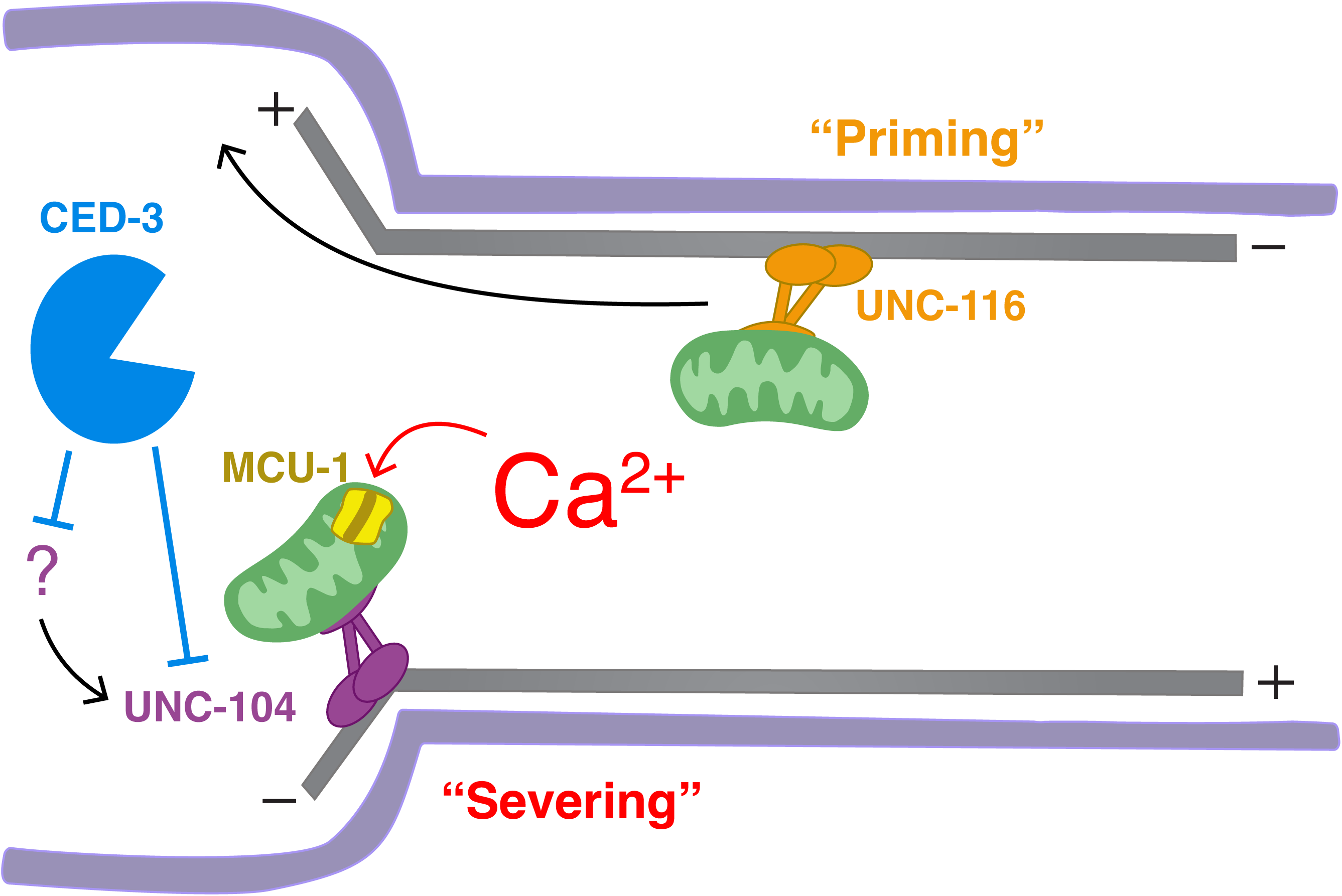
Model for regulation of CCE by mitochondrial transport. UNC-116/Kinesin 1 and UNC-104/Kinesin 3 play opposing roles in transporting mitochondria in the context of CCE. UNC-116/Kinesin 1 transports mitochondria retrogradely into the soma to prime the cell for CCE. CED-3/Caspase ensures mitochondria remain remote from the soma-process junction and in the soma by negatively regulating UNC-104/Kinesin 3. This permits localized Ca^2+^ spiking at the soma-process junction cytosol and some-process junction severing as the first event of CCE. In the absence of CED-3/Caspase, UNC-104/Kinesin 3 performs a non-canonical function of short-range mitochondrial anterograde transport to the soma-process junction. Mitochondria so-transported take up cytosolic Ca^2+^ via the uniporter MCU-1.

## Discussion

In this study, we have taken an *in vivo* genetic and cell biological approach to demonstrate that mitochondrial transport and mitochondrial Ca^2+^ uptake are important in the regulation of *C. elegans* Compartmentalized Cell Elimination (CCE), and thus provide further molecular insights into the regulation of specialized cell death and neurite pruning. Our study highlights that specialized cell elimination can be regulated at specific cellular microdomains, with Ca^2+^ spikes, caspase activity and mitochondrial transport events taking place at a dedicated severing point.

Mitochondria have long been implicated in neuroprotection. Elevated mitochondrial activity can rescue neurodegeneration in disease models in mice (47). In addition, work in primary cerebrocortical neurons and mice suggests that mitochondria in distal neuronal compartments protect against in neuron retraction (such as synaptic pruning or axon and dendrite regression) and local caspase activation(48). Consistent with this, previous work in *C. elegans* has shown that, following injury, mitochondria can protect axons from degeneration(40) and can aid in regeneration(15). To accomplish this, mitochondria are transported to the relevant area. In *C. elegans*, UNC-116/Kinesin 1 has been shown to play an important role in this transport(40) and is also important for maintenance of mitochondrial density in motor(40) and touch receptor (49) neurons.

Kinesin 1 is known to be involved in anterograde mitochondrial transport(50). Here we show that UNC-116/Kinesin 1, in the polarized but non-neuronal TSC, instead of transporting mitochondria to energy-requiring sites anterogradely, retrogradely transports mitochondria away from sites destined for elimination prior to morphological indications of cell death initiation. Thus, in the context of programmed cell elimination, UNC-116/Kinesin 1 serves in cell death priming through region-specific mitochondrial exclusion.

In addition to UNC-116/Kinesin 1, we present as a versatile motor the founding member of the kinesin-3 family, UNC-104/Kinesin 3(51). We identify mitochondria as a non-canonical cargo. Pioneering studies in the nematode first showed UNC-104/Kinesin 3 to transport pre-synaptic vesicles (26). Subsequent studies from the murine brain showed Kinesin 3/KIF1A as a neuron-specific member of the kinesin motor superfamily functioning in fast anterograde axonal transport of synaptic vesicle precursors(52). Interestingly, the kinesin-3 family domain sequence is highly conserved, with a motor domain composed of an ATP catalytic center, a microtubule-binding surface, and a pleckstrin homology (PH) region. The PH region acts as a lipid-binding domain for cargo(53). Structure-function studies of UNC-104/Kinesin 3’s PH domain further may provide insights into potential cargo adaptability of this kinesin family. Interestingly, while *C. elegans* UNC-104/Kinesin 3 itself, to our knowledge, has not been previously demonstrated to carry mitochondria, there is in fact precedent for other kinesin-3 family members to do so (54), including KIF1B in mouse Neuro2a cells(54), Nkin2 / Nkin3 in *Neurospora crassa*(55) and KLP-6(56) in *C. elegans.* Our report that UNC-104/Kinesin 3 can have a function beyond transport of synaptic vesicle cargo is further substantiated by our observation that UNC-104/Kinesin 3 is expressed in a non-neuronal cell and important for its programmed fate. Additionally, in our model, UNC-104/Kinesin 3 appears to serve in directed, but short-range, transport, while kinesins as a family are known to have processive motility, carrying organelles over large distances(57).

In neurons, both synaptic vesicles and mitochondria need to be transported along the neurite over long distances, yet the nature of these cargo is vastly different. This may be accommodated by a diverse repertoire of motors, adaptors and modifications. In terms of size, synaptic vesicles are of the order of 40nm whereas mitochondria can range from 0.5-1µm. A question of long-standing interest is how kinesins can display long-range transport along microtubules without being disrupted by steric hindrance(58). It is possible that UNC-116/Kinesin 1, a bonafide mitochondrial motor, is structurally equipped to perform longer range transport of a larger cargo whereas TSC UNC-104/Kinesin 3 displays short-range dynamics as it carries mitochondria as a larger cargo than for which it was designed. How UNC-104/Kinesin 3 associates with mitochondria remains an open question for future studies. It is possible it acts in conjunction with an adaptor specific to the TSC.

We also find that UNC-116/Kinesin 1 and UNC-104/Kinesin 3-dependent transport of cell-preserving mitochondria impact CCE in different ways based on the molecular context and cellular subdomain. In an otherwise wild-type background, loss of UNC-116/Kinesin 1 only results in a distal process elimination defect, with the rest of the cell being removed. This distal remnant always harbors mitochondria, suggesting that distally arrested mitochondria protect this specific domain of the cell from elimination. Following failure to sever, the TSC reverts to its earlier intact morphology in absence of CED-3/Caspase. We thus propose that the severing event is essential for CCE completion, and the protective impact of mitochondria depends on which cell compartment the mitochondria are located: distal mitochondria can only protect the cell locally and severing point mitochondria can save the entire cell. Whether this is a function of the subcellular architecture of that domain and its relative importance to overall cell structure remains to be determined.

One interesting observation is that the proximal part of the TSC process appears to show mixed microtubule polarity, while plus-end-in distally, suggesting compartment-specific specialization at the level of the cytoskeleton that may dictate the importance of that domain to CCE and how its dismantling is regulated. In addition, we note that mitochondria are protective for the TSC process and soma-process junction, but not the soma. This may be due to fundamental differences between these cell compartments morphologically and molecularly and how they respond to mitochondrial function.

This point motivates the question of how mitochondria are transported to and arrested to function at specific cellular microdomains at the appropriate time. Well-established work has addressed this in neurons. Axonal mitochondria are known to be recruited to synaptic terminals through adaptors such as syntaphilin (SNPH) which mediates immobilization of axonal mitochondria through binding to KIF5 in response to changes in neuronal activity and sensing mitochondrial Rho guanosine triphosphatase-Ca^2+^ (59). Additionally, Ca^2+^ signaling can regulate both anterograde and retrograde mitochondrial movement. For example, the small GTPase Miro plays an important role in neurons in the resistance to excitotoxicity by promoting the arrest of mitochondrial motion by binding to Ca^2+^ allowing it to directly interact with the motor domain of Kinesin 1, preventing the latter from further interacting with microtubules(60). Our work expands on these concepts in a new context, cell death, by implicating local Ca^2+^ spikes as a potential cue for UNC-104/Kinesin 3 to carry mitochondria anterogradely to the TSC severing site.

In addition to suggesting Ca^2+^ as a possible transport cue, we also propose it as required for specialized cell elimination. We show mitochondria can serve to be protective to the cell in a compartment-specific manner by virtue of their Ca^2+^ buffering capacity. Ca^2+^ transients have long been appreciated to modulate varying forms of cell death, including apoptosis, necrosis and more recently other forms such as autophagic cell death and anoikis(61). Ca^2+^ transients in the cytoplasm are generally important for neuronal function and mitochondria are especially critical for Ca^2+^ homeostasis. Mitochondrial Ca^2+^ overload can lead excitotoxicity and neuronal death(62). The mitochondrial Ca^2+^ uniporter (MCU) was discovered to uptake Ca^2+^ through the inner mitochondrial membrane following stress(63). In *Drosophila* models of neurodegenerative diseases, mitochondria have been shown to exhibit Ca^2+^ buffering capacity that is regulated by MCU (64). Interestingly, it has also been shown in the non-neuronal context of the adult *C. elegans* epidermis(65), that MCU-1 is required for rapid mitochondrial Ca^2+^ uptake following wounding. This in turn leads to mitochondrial ROS production which promotes cytoskeletal responses that facilitate wound closure(45). An important facet of tissue regeneration is the rapid initiation of wound repair and wounding triggers intracellular Ca^2+^ elevation as a damage signal(66–68). It will be interesting to explore CCE as a programmed injury model and the role of ROS in the absence of CED-3/Caspase to prevent severing and mechanistic overlaps between regeneration and wound healing with the preservation of the soma-process junction during CCE. Mitochondrial Ca^2+^ uptake at the soma-process junction may either prevent the completion of severing or allows for speedy repair.

In *Drosophila* sensory neurons, it has been shown that pruning can be temporally and spatially triggered by compartmentalized Ca^2+^ signaling at select dendritic branches(69). Work on Drosophila commissural axons implicate a cell-intrinsic mechanism controlling compartmentalized receptor localization (70–73). In *C. elegans*, specific areas of axons of a subset of interneurons exhibit localized Ca^2+^ dynamics regulating behavior(13, 74–76). While several studies have shown that mitochondria can be transported to sites for protection and implicate Ca^2+^, our work, in revealing a specific role for localized Ca^2+^ uptake in specialized cell elimination, provides additional insights on developmental pruning.

In our CCE model, local Ca^2+^ uptake at the soma-process junction is performed by mitochondria transported to this region by UNC-104/Kinesin 3 to protect the cell from death in the absence of CED-3/Caspase. Our genetics and the observation that UNC-104/Kinesin 3 protein is only seen in the TSC prior to CCE onset in the absence of CED-3/Caspase suggests CED-3 activity negatively regulates UNC-104/Kinesin 3. Notably, caspase activity in wild-type and the UNC-104/Kinesin 3 signal in *ced-3* mutants spatially overlap, both seen in the soma-process junction. Whether CED-3/Caspase targets UNC-104/Kinesin 3 for cleavage directly or a partner or adaptor remains to be examined.

Unexpectedly, our data show that in the absence of CED-3/Caspase, other than complete soma-process detachment, stereotyped morphological feature of CCE are seen. The only event that remains incomplete appears to be severing. This would presumably be due to the function of UNC-104/Kinesin 3 and its ability to transport mitochondria to the severing site. We propose cytosolic Ca^2+^ spikes are important for severing during CCE and our data do support the idea that UNC-104/Kinesin 3 and mitochondrial Ca^2+^ uptake prevent these localized spikes. However, when UNC-104/Kinesin 3 or MCU-1 are removed, while the Ca^2+^ signal does indeed return, in the absence of CED-3/Caspase, the severing event still fails. As such, while our data supports a model for UNC-104/Kinesin 3 and mitochondrial Ca^2+^ uptake being important for Ca^2+^ signaling at the soma-process junction, in the absence of CED-3/Caspase, the precise role of this Ca^2+^ in CCE is yet to be determined and is an important and exciting future direction. Notably, CED-3/Caspase is clearly of very fundamental importance to CCE, as in the case of apoptosis, and may be playing several roles to ensure this complex program takes place properly such that loss of this protease has many consequences. Regulating mitochondrial transport is likely just one of many roles CED-3/Caspase plays, and its loss may deregulate a number of events required for CCE completion. This implicates CCE as a rich setting for studying additional cell death factors regulated by CED-3/Caspase directly or indirectly and additional specialized cell death events.

We have previously proposed that CCE can serve as a model for pruning, which is also caspase dependent, based on morphological similarities and its occurrence in a set of nematode sex-specific sensory neurons(35, 77). It has been shown in cultured neurons that caspase-3 induction by photoactivation in specific dendritic regions leads to dendritic retraction and local elimination of spines, but not cell death, suggesting spatial restriction of caspase activity(78). Our prior work has also suggested that CED-3/caspase acts independently in the different TSC sub-compartments based on the range of compartment-specific defects of the *ced-3* hypomorph(35). We find that localized caspase activity is important for the completion CCE, with caspase activity seen at the severing site, but not distally, in wild-type animals, leading to pruning like events. An unexpected observation in our current study is that CED-3/Caspase appears not to be required for the very initiation of CCE events, namely for the attempt at severing and for the appearance of other CCE hallmarks across compartments, but rather the completion of severing and soma-process detachment. Thus, we describe a caspase function at a very specific spatial and temporal resolution. This may be solely due to the localized nature of CED-3 activity, as indicated by the caspase activity sensor used in this study. As such, the more pertinent question would be: what potentiates region-specific caspase activity? There is evidence of region-specific pruning being regulated by caspase gradients(4). The observation that the completion of soma-process severing appears to require CED-3/Caspase and Ca^2+^ also begs the question of what the exact caspase, Ca^2+^ dependent and region-specific effector severs this cellular subdomain so specifically. A plausible explanation is that these effectors would target components of the cytoskeleton.

Spatially, the severing point at the TSC soma-process junction is reminiscent of the AIS of neurons. The AIS cytoskeleton has been shown to undergo rapid and irreversible proteolysis following neuronal injury, without cell death or axonal destruction, and that this requires the Ca^2+^-dependent cysteine protease calpain(79, 80). Interestingly, the AIS has been shown to be important for axon regeneration and functional recovery following injury(81). Our study, in underscoring the importance of specific compartments of a specialized cell to its elimination, opens the question of the specific cell regressive contribution of the AIS to neuronal death, neurite pruning and caspase function. Additionally, whether mitochondria play specialized roles in the AIS would be an interesting avenue to explore motivated by this work.

Neurons and other morphologically complex cells are proposed to share compartmentalization mechanisms(16). Our work, by highlighting new facets of specialized cell elimination during development and implicating mitochondrial transport and Ca^2+^ uptake in its regulation, positions the study of CCE as a valuable setting to better understand the fundamentals of neurite pruning and neurodegeneration.

---

**Supplemental Figure S1. Mitochondrial position relative to soma-process junction in WT and ced-3(n717). (A-E)** Wild-type (F-J) *ced-3(n717)*.

## Supplemental Files

### Supplemental Movies and Stacks

**Video S1.** TSC WT mitochondria timelapse, related to Figure 2.

**Video S2.** TSC WT mitochondria timelapse, related to Figure 3.

**Video S3.** TSC *ced-3(n717)* mitochondria timelapse embryo, related to Figure 3.

**Video S4.** L1 TSC *ced-3(n717)* EBP-2::GFP Kymograph Movie (Proximal), related to Figure 4.

**Video S5.** L1 TSC *ced-3(n717)* EBP-2::GFP Kymograph Movies (Distal), related to Figure 4.

**Video S6.** UNC-116 localization in WT 1.5-fold embryo, related to Figure 1.

**Video S7-S12.** TSC mitochondria WT embryo, related to Figure 2.

**Video S13-S16.** TSC GFP::C3Ai WT embryo, related to Figure 3.

**Video S17-S20.** TSC GFP::C3Ai *ced-3(n717)* embryo, related to Figure 3.

**Video S21-S24.** UNC-104::GFP WT embryo, related to Figure 4.

**Video S25-S28.** UNC-104::GFP *ced-3(n717)* embryo, related to Figure 4.

**Video S29, S30.** TSC mitochondria *ced-3(n717); unc-104(e1265)* embryo, related to Figure

**Video S31-S33.** TSC GCaMP5a WT embryo, related to Figure 5.

**Video S34-S36.** TSC GCaMP5a *ced-3(n717)* embryo, related to Figure 5.

**Video S37-S39.** TSC GCaMP5a *ced-3(n717); unc-104(e1265)* embryo, related to Figure 5.

**Video S40-S42.** TSC GCaMP5a *ced-3(n717); mcu-1(ju1154)* embryo, related to Figure 5.

### Supplemental Tables

**Supplemental Table 1.** Plasmids used in this study

**Supplemental Table 2.** Transgenes used in this study

## Materials and Methods

### C. elegans methods

*C. elegans* strains were cultured per standard methods and were grown at 20°C(82). Wild-type animals were of the Bristol N2 subspecies. For most tail-spike cell (TSC) experiments, one of three integrated reporters were used: *nsIs435*, *nsIs431,* or *nsIs686*. Integration of extrachromosomal arrays was performed using UV and trioxsalen (Sigma T2137). Other transgenic strains generated are described in **Supplemental Table 2**. Larvae were scored at 20°C at the L1 stage.

### Mutants

The list of mutant *C. elegans* strains used in this study is as follows:

LGI-
LGII-*unc-104(e1265)*
LGIII-*unc-116(ns827), unc-116(e2310), ced-4(n1162), clk-1(qm30)*
LGIV-*ced-3(n717), ced-3(n2427), mcu-1(ju1154)*
LGV-
LGX-*dhc-1(js319)*

### Germline transformation and rescue experiments

Germline transformation was carried out as previously described(83). All plasmids were injected at between 1 and 20ng per µl. pBSK was used to adjust the DNA concentration of injection mixtures if necessary. All rescue experiments were done with *myo-2*p::GFP as a co-injection marker along with *cdh-3*p::mCherry to label the TSC.

### Primers and plasmid construction

Plasmids were generated by either Gibson Cloning or restriction digest. Primer sequences and information on the construction of plasmids used in this study are provided in **Supplemental Table 1.**

### Transgenes

The full list of transgenes is described in **Supplemental Table 2.** The full length or fragment of the *aff-1* promoter was used to label the TSC.

### CRISPR Cas9 genome editing

To generate TSC404, GFP was introduced to the C-terminus of *unc-104* via CRISPR/Cas9 using the crRNA-5’-GCAATTGAAGATGATGATGT-3’. To generate TSC459, GFP was introduced to the C-terminus of *unc-116* via CRISPR/Cas9 using the crRNA 5’-GTTAAATTTGATAATACGGT-3’. Mutants were generated using a co-injection strategy(84). Guide crRNA, repair single-stranded DNA oligos, tracrRNA, and buffers were ordered from IDT. The strain PHX6842, *mcu-1(syb6842); ced-3(n2427)*; *nsIs435* was made by deleting base pairs 468 to 1364 from the gene *mcu-1* in the background of *ced-3(n2427)*; *nsIs435* and was generated by Suny Biotech (Suzhou, Jiangsu, China 215028).

### Scoring of TSC/CCE defects

TSC death was scored at the L1 stage. Animals were synchronized allowing gravid hermaphrodites to lay eggs overnight, washing off larvae the next day and waiting 5 hours for hatched L1s. L1s synchronized were then mounted on slides on 2% agarose-water pads, anaesthetized in 10 mM sodium azide and examined on a wide-field Axioscope A1 (Zeiss) at 40X magnification (NA 1.3). The TSC was identified by fluorescence (from reporter transgenes) as well as by its location and morphology.

### Mutagenesis and mutant identification

*nsIs435* animals were mutagenized using 75 mM ethyl methanesulfonate (M0880, Sigma) for 4 hours at 20 °C. Approximately 27,000 F2 progeny were screened for TSC persistence on a Zeiss Axio-Scope A1 at 40X. *unc-116 (ns827)* was identified from analysis of SNP-mapping(85), Whole Genome Sequencing data, fosmid rescue and candidate gene analysis.

### Microscopy and image processing

Images were collected on a Nikon TI2-E Inverted microscope using a CFI60 Plan Apochromat Lambda 60x Oil Immersion Objective Lens, N.A. 1.4 (Nikon) and a Yokogawa W1 Dual Cam Spinning Disk Confocal. Images were acquired using NIS-Elements Advanced Research Package. For still embryo imaging, embryos were anesthetized using 0.5 M sodium azide. Larvae were paralyzed with 10mM sodium azide.

Light sheet imaging was performed using a customized ASI diSPIM(86) equipped with symmetric 40X Nikon CFI APO NIR objectives and imaged onto Hamamatsu Orca Fusion (C14440-20UP) cameras equipped with Hamamatsu W-View Gemini channel splitters for simultaneous two-color imaging. Imaging was performed with a single view 1 micron z-spacing and volumes were acquired every 3 minutes using the diSPIM control plug-in for Micro-Manager 1.4(87) and a uniform background subtraction of 100 counts was applied using ImageJ.

EBP-2 imaging was done on *ced-3(n717)* mutant L1 larvae following previously published protocol(88) on the Nikon TI2-E Inverted microscope above. Kymographs were generated using ImageJ.

### Quantification of mitochondria

Length of TSC process and number of mitochondria in the TSC process was counted using ImageJ using analyze particle tool. Mitochondria number/TSC process length was calculated in Excel and these values were input in GraphPad to generate graph.

### Statistics and reproducibility

The sample sizes and statistical tests were selected based on previous studies with similar methodologies. Sample sizes were not determined using statistical methods. All experiments were repeated at least two to three times, as indicated, giving similar results. Independent transgenic lines were treated as independent experiments. Quantification of TSC persistence was done using an unpaired two-tailed *t*-test (GraphPad). For all figures, mean ± standard error of the mean (s.e.m.) is represented.

## Acknowledgements

Some strains were provided by the CGC, which is funded by NIH Office of Research Infrastructure Programs (P40 OD010440). PG is funded by a National Institutes of Health-National Institute of General Medical Sciences Maximizing Investigators’ Research Award (MIRA) (R35GM142489) and is a Cancer Prevention Research Institute of Texas (CPRIT) Scholar in Cancer Research (RR100091) This work was also supported by NIH MIRA grants R35GM151199 to PKS, R35GM128885 to MWP, a CPRIT Scholar in Cancer Research (RR160053), and NIH grants R01HD103610 and R35NS105094 to SS. We thank the Ghose Lab for comments on the manuscript.

## Author Contributions

RS and PG designed the experiments and wrote the paper. RS, SP, AE, PG, PKS performed experiments and analyzed data. GC provided significant technical assistance. PG and RS conceptualized the study, with early contributions from MWP and SS and early resources from SS.

## Competing Interests

The authors declare no competing interests.

## Notes

### Competing Interest Statement

The authors have declared no competing interest.

