## Supplementary figures and images for "Mitochondria transported by Kinesin 3 prevent localized calcium spiking to inhibit caspase-dependent specialized cell death"

### Supplemental Figure 1

#### Sharmin et. al, Supplemental FIGURE S1

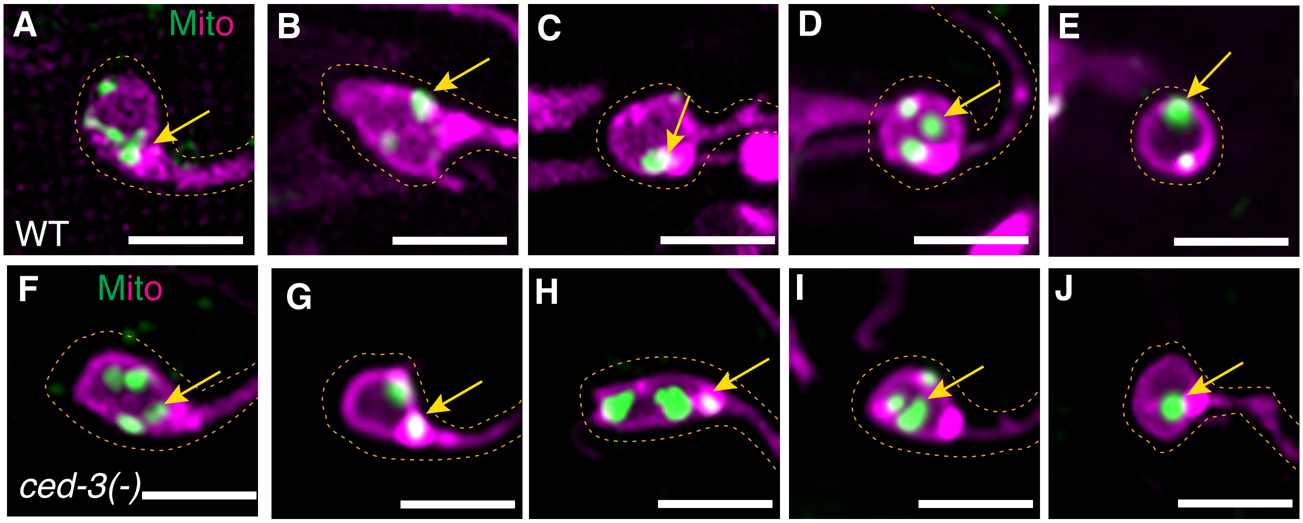
