## Supplemental Table 1 for "Mitochondria transported by Kinesin 3 prevent localized calcium spiking to inhibit caspase-dependent specialized cell death"

### Supplemental Table 1. Plasmids used in this study

| Gibson Cloning | | | | | | | |
|---|---|---|---|---|---|---|---|
| **Plasmid name** | **Plasmid Description** | **Insert Forward Primer** | **Insert Reverse Primer** | **Backbone Forward Primer** | **Backbone Reverse Primer** | **Insert Template** | **Backbone Template** |
| pPG294 | TSCp::unc-104 cDNA (cell specific rescue) | tctagaggatcccgggattggccaaag | GGCGCGCCgttccgaatattatgaaatg | tccatactttctcatttcataatattcggaacGGCGCGCCaaaATGTCATCGGTTAAAGTAGCTGTACG | tacctttgggtcctttggccaatcccggggatcctctagaTTATGAAGCAGCAATTGAAGATGATGATG | | pSM::SL2::mCherry |
| pPG331 | TSCp::unc-104GFP_tomm7 (mito rescue) | tccatactttctcatttcataatattcggaacGGCGCGCCaaaATGTCATCGGTTAAAGTAGCTGTACG | TACTgcttccgccggtacctccactgccaccgctagtactTGAAGCAGCAATTGAAGATGATGATGTTG | agtactagcggtggcagtggaggtac | GGCGCGCCgttccgaatattatgaaatg | unc-104 cDNA | pPG174 (below) |
| pPG445 | TSCp::mcu-1 cDNA (cell specific rescue) | tccatactttctcatttcataatattcggaacGGCGCGCCaaaATGAGGAATGGCCGATGCTTGGTG | tacctttgggtcctttggccaatcccggggatcctctagaTTACTTTTCAGCTTCCAAATTGGATAAATAG | GGCGCGCCgttccgaatattatg | tctagaggatccccgggattggc | mcu-1 cDNA | pSM::SL2::mCherry |
| pPG245 | unc-116p::mKate2 (expression) | GATCCTCTAGAGTCGACCTGCAGGC | GCTTCATATGCATGTTTTCgTTAATGAGCTCGGAGACCATtttcagagaataatgatcaccagttggc | ATGGTCTCCGAGCTCATTAAcGAAAAC | GATCCTCTAGAGTCGACCTGCAGGC | unc-116 genomic DNA | pPG146=cdh-3p::mKate2 |
| pPG246 | TSCp::unc-116 cDNA (cell specific rescue) | tccatactttctcatttcataatattcggaacGGCGCGCCaaaATGGAGCCGCGGACAGACGGAG | Tacctttgggtcctttggccaatcccggggatcctctagacgatgagttgatgttgttgttgggagac | | | | pSM::SL2::mCherry |
| pPG401 | TSCp::UNC-104::mCherry | ATTGAAATCTCCAACATCATCATTCAATTG | caagttggtaatggtagcgaccggcgctcagttgGAATTCT | ctctattctcacaaaaaacatatg | ggagaaagagcatgtagg | pPG331 (above) | pPG270=TSCp::lnp-1 cDNA_SL2::mCherry |

| | | | | | | | |
|---|---|---|---|---|---|---|---|
| | | CTGCTTCAGGAGCATCGGGAGCCTCAGGAGCATCGATGGTGagcaagggcgaggaggataaca | TActtgtacagctcgtccatgccgc | | | | (unpublished) |
| pPG274 | TSCp::EBP-2::GFP | tccatactttctcatttcataatattcggaacGGCGCGCCttcagaaaccgaggcaaaATGGTCG | caagttggtaatggtagcgaccggcgctcagttgGAATTCctatttgtatagttcatccatgccatgt | GGCGCGCCgttccgaatattatg | GAATTCcaactgagcgccggtc | Pklp-6::epb-2::gfp (A gift grom Maureen Barr) | pPG270 |
| **Restriction Digest Based Cloning** | | | | | | | |
| **Plasmid name** | **Plasmid Description** | **Insert Forward Primer** | **Insert Reverse Primer** | **Vector** | | **Insert Template** | |
| pPG174 | TSCp::unc116GFP_tomm7 | cccccGGCCGGCCctattgacgataaccaccttacaaac | ccaatcccggggatcctctagaGGCGCGCCggggg | Punc-47::unc-116_GFP::tomm-7 | | aff-1 promoter 1393 bp (Fse-1, AscI) | |
| pPG115 | TSCp::GCaMP5a_SL2_myrmcherry | ccccccGGCCGGCCctattgacgataaccaccttacaaacacc | ccccccGGCGCGCCgttccgaatattatgaaatgagaaagtatgG | pSM::SL2::mCherry | | GCaMP5a synthesized | |
| pPG113 | TSCp::GFP::C3Ai_SL2::myrmCherry | ccccccGGCGCGCCaaaATGATCAAGATTGCCACACGGAAATAT | ggggggGTTAACTTAATTCGGCAGATTGTCGACGCGCAT | pSM::SL2::mCherry | | GFP::C3Ai synthesized | |
| pPG112 | TSCp::mitoGFP | ccccccGGCGCGCCaaaATGGCACTCCTGCAATCACGTCTCC | ggggggGTCGACTTTGTATAGTTCATCCATGCCATGTGTAATC | pglr-1::mitoGFP (Ghose et al, 2013) | | aff-1 promoter 1391 bp (AscI, SalI) | |
| pPG114 | TSCp::mitoGFP_SL2_myrmcherry | cccccggcgcgccaaaATGGCACTCCTGCAATCACGTC | ccccccagttaactaggtgaaagtaggatgagacaggatatc | pSM::SL2::mCherry | | pPG112 (AscI, EcoRV) | |
