## Supplemental Table 2 for "Mitochondria transported by Kinesin 3 prevent localized calcium spiking to inhibit caspase-dependent specialized cell death"

**Supplement Table 2: List of transgenes and strains**

| Strain | Genotype | Comments |
|---|---|---|
| TSC6 | *unc-116(e2310); nsIs435* | *nsIs435=aff-1*p::myrGFP |
| TSC16 | *ced-3(n717); nsIs435; mccEx006* | *mccEx006*=pPG245; *nsIs435=aff-1*p::myrGFP |
| TSC17 | *ced-3(n717); nsIs435; mccEx007* | *mccEx007*=pPG245; *nsIs435=aff-1*p::myrGFP |
| TSC32 | *unc-116(ns827); mccEx019* | *mccEx019*=pPG246; *nsIs435=aff-1*p::myrGFP |
| TSC33 | *unc-116(ns827); mccEx020* | *mccEx020*=pPG246; *nsIs435=aff-1*p::myrGFP |
| TSC40 | *unc-116(ns827); nsls686; mccIs005* | *mccIs005*=pPG112; *nsIs686=aff-1*p::mKate2 |
| TSC66 | *ced-3(n717); nsls686; mccIs005* | *mccIs005*=pPG112; *nsIs686=aff-1*p::mKate2 |
| TSC74 | *unc-104(e1265); ced-3(n2427); nsIs435* | *nsIs435=aff-1*p::myrGFP |
| TSC89 | *unc-104(e1265); nsIs435* | *nsIs435=aff-1*p::myrGFP |
| TSC90 | *trak-1(tm1572); ced-3(n2427); nsIs435* | *nsIs435=aff-1*p::myrGFP |
| TSC91 | *trak-1(tm1572); nsIs435* | *nsIs435=aff-1*p::myrGFP |
| TSC96 | *unc-116(ns827);ced-3(n2427);nsIs435* | *nsIs435=aff-1*p::myrGFP |
| TSC98 | *unc-116(ns827); nsIs435; mccEx047* | *mccEx047*=pPG174; *nsIs435=aff-1*p::myrGFP |
| TSC99 | *unc-116(ns827); nsIs435; mccEx048* | *mccEx048*=pPG174; *nsIs435=aff-1*p::myrGFP |
| TSC100 | *unc-116(ns827); nsIs435; mccEx049* | *mccEx049*=pPG174; *nsIs435=aff-1*p::myrGFP |
| TSC105 | *unc-104(e1265); ced-3(n2427); nsIs435; mccEx053* | *mccEx053*=pPG294; *nsIs435=aff-1*p::myrGFP |
| TSC106 | *unc-104(e1265); ced-3(n2427); nsIs435; mccEx054* | *mccEx054*=pPG294; *nsIs435=aff-1*p::myrGFP |
| TSC107 | *unc-104(e1265); ced-3(n2427); nsIs435; mccEx055* | *mccEx055*=pPG294; *nsIs435=aff-1*p::myrGFP |
| TSC260 | *ced-3(n717); nsIs532* | *nsIs532*=pPG114 |
| TSC261 | *ced-3(n717); nsIs560* | *nsIs560*=pPG113 |
| TSC264 | *unc-116(ns827); ced-3(n717); nsls686; mccIs005* | *mccIs005*=pPG112; *nsIs686=aff-1*p::mKate2 |
| TSC277 | *ced-3(n2427); dhc-1(js319); nsIs435* | *nsIs435=aff-1*p::myrGFP |

| | | |
|---|---|---|
| TSC304 | *unc-104(e1265); ced-3(n2427); nsIs435; mccEx156* | *mccEx156*=pPG331; *nsIs435=aff-1*p::myrGFP |
| TSC305 | *unc-104(e1265); ced-3(n2427); nsIs435; mccEx157* | *mccEx157*=pPG331; *nsIs435=aff-1*p::myrGFP |
| TSC306 | *unc-104(e1265); ced-3(n2427); nsIs435; mccEx158* | *mccEx158*=pPG331; *nsIs435=aff-1*p::myrGFP |
| TSC386 | *unc-116(ns827); ced-3(n717); nsls686; mccIs005; mcc27* | *mcc27*=Small deletion few nucleotides upstream PAM site; *mccIs005*=pPG112; *nsIs686=aff-1*p::mKate2 |
| TSC404 | *unc-104(e1265); mcc28; ced-3(n717)* | *mcc28*=GFP inserted just before stop codon (C terminus) |
| TSC405 | *unc-104(e1265); mcc28; ced-3(n717); nsIs686* | *mcc28*=GFP inserted just before stop codon (C terminus); *nsIs686=aff-1*p::mKate2 |
| TSC407 | *ced-4(n1162); nsIs686; mccIs015* | *mccIs015*=pPG274; *nsIs686=aff-1*p::mKate2 |
| TSC409 | *clk-1(qm30); ced-3(n2427); nsIs435* | *nsIs435=aff-1*p::myrGFP |
| TSC459 | *unc-116(mcc33)* | *mcc33*=[UNC-116::GFP] N2; CRISPR mutant, GFP insertion after UNC-116 |
| TSC472 | *clk-1(qm30); nsIs435* | *nsIs435=aff-1*p::myrGFP |
| TSC489 | *N2; mcc33; mccEx207* | *mcc33*=[UNC-116::GFP] N2; CRISPR mutant, GFP insertion after UNC-116; *mccEx207=aff-1*p::myrmCherry |
| TSC514 | *ced-3(n717); mccIs002; mccIs050* | *mccIs002*=pPG112; *mccIs050*=pPG401 |
| TSC521 | *dhc-1(js319); nsIs435* | *nsIs435=aff-1*p::myrGFP |
| TSC522 | *mcu-1(ju1154); nsIs435* | *nsIs435=aff-1*p::myrGFP |
| TSC577 | *N2; mcc28; mccIs094* | *mcc28*=UNC-104::GFP inserted just before stop codon (C terminus); *mccIs094=aff-1*p::myrmCherry |
| TSC580 | *ced-3(n717); mcc28; mccIs100* | *mcc28*=UNC-104::GFP inserted just before stop codon (C terminus); *mccIs100=aff-1*p::myrmCherry |
| TSC581 | *N2; mcc33; mccIs094* | *mcc33*=[UNC-116::GFP] N2; CRISPR mutant, GFP insertion after UNC-116; *mccIs094=aff-1*p::myrmCherry |
| TSC582 | *N2; mcc33; mccIs100* | *mcc33*=[UNC-116::GFP] N2; CRISPR mutant, GFP insertion after UNC-116; *mccIs100=aff-*1p::myrmCherry |
| TSC681 | *ced-(n717); nsIs560* | *nsIs560*=pPG113 |
| TSC682 | *ced-(n717); mcu-1(ju1154); nsIs560* | *nsIs560*=pPG113 |
| TSC683 | *ced-(n717); unc-104(e1265); nsIs560* | *nsIs560*=pPG113 |
| TSC685 | *mcu-1(syb6842); ced-3(n2427); nsIs435; mccEx276* | *mccEx276*=pPG445; *nsIs435=aff-1*p::myrGFP |

| | | |
|---|---|---|
| TSC686 | *mcu-1(syb6842); ced-3(n2427); nsIs435; mccEx277* | *mccEx277*=pPG445; *nsIs435*=*aff-1*p::myrGFP |
| TSC688 | *mcu-1(syb6842); ced-3(n2427); nsIs435; mccEx279* | *mccEx279*=pPG445; *nsIs435*=*aff-1*p::myrGFP |
| TSC693 | *ced-3(n717); unc-104(e1265); nsIs686; mccIs005* | *mccIs005*=pPG112; *nsIs686*=*aff-1*p::mKate2 |
| OS8095 | *ced-3(n717); nsIs435* | *nsIs435*=*aff-1*p::myrGFP |
| OS9235 | *ced-3(n2427); nsIs435* | *nsIs435*=*aff-1*p::myrGFP |
| OS9985 | *N2; nsIs532* | *nsIs532*=pPG114 |
| OS10204 | *N2; nsIs560* | *nsIs560*=pPG113 |
| OS11918 | *unc-116(ns827); nsIs435* | *nsIs435*=*aff-1*p::myrGFP |
| PHX6842 | *mcu-1(syb6842); ced-3(n2427)* | |
